# Spatial Awareness of a Bacterial Swarm

**DOI:** 10.1101/341529

**Authors:** Harshitha S. Kotian, Shalini Harkar, Shubham Joge, Ayushi Mishra, Amith Zafal, Varsha Singh, Manoj M. Varma

## Abstract

Bacteria are perhaps the simplest living systems capable of complex behaviour involving sensing and coherent, collective behaviour an example of which is the phenomena of swarming on agar surfaces. Two fundamental questions in bacterial swarming is how the information gathered by individual members of the swarm is shared across the swarm leading to coordinated swarm behaviour and what specific advantages does membership of the swarm provide its members in learning about their environment. In this article, we show a remarkable example of the collective advantage of a bacterial swarm which enables it to sense inert obstacles along its path. Agent based computational model of swarming revealed that independent individual behaviour in response to a two-component signalling mechanism could produce such behaviour. This is striking because independent individual behaviour without any explicit communication between agents was found to be sufficient for the swarm to effectively compute the gradient of signalling molecule concentration across the swarm and respond to it.

## Introduction

Bacteria are perhaps the simplest living systems capable of complex behaviour involving sensing and coherent, collective behaviour an example of which is the phenomena of swarming, where bacteria colonize a solid surface (typically, nutrient loaded agar) in geometric patterns characteristic of different species (Ben-Jacob, 1997; Kearns, 2010). A natural and fascinating question in this regard is how the information gathered by individual members of the swarm through sensing their respective local environments is shared across the swarm leading to coordinated swarm behaviour. Specifically, does membership of the swarm provide its members any advantage in learning more about their environment? For instance, are they able to extract more information about their surroundings collectively than acting alone? Swarming involves several, possibly collective, decision making steps such as quorum sensing (Daniels et al., 2004). The swarming patterns produced by the Gram-negative bacteria *Pseudomonas aeruginosa* (PA) are special due to the presence of long straight segments (tendrils) in its swarming pattern [Figure 1 a)]. The swarming pattern produced by PA is unique in two major aspects. Firstly, the typical tendril length (^~^ cm) is nearly 4 orders of magnitude compared to the micron-sized individual swarm member. Considering the large stochastic fluctuations likely to be experienced by a micron sized individual member, it is remarkable that collective motion of these individuals produces such long straight segments. The long directional persistence leading to this highly anisotropic collective motion is worthy of detailed mathematical modelling. Indeed, there have been several attempts to model the swarming pattern of PA based on a highly mechanistic multiscale model(Du et al., 2011), as a population dispersal phenomenon using spatial kernels(Deng et al., 2014) and based on Marangoni forces(Du et al., 2012). However, so far none of these have succeeded in producing patterns with similar branching statistics similar to the one seen in Figure 1 a) [SI Section1 Figure S2]. While generic models such as the Vicsek model [(Vicsek et al., 1995)] can explain large directional persistence collections of stochastically moving particles, one would like to have a more detailed model involving experimentally accessible parameters of the system which produces statistically identical patterns as seen in the experimental system. While this is an open problem, in this paper we focus on a different shortcoming of the existing models which is that these models have primarily been employed to study the pattern formation and do not describe other aspects of swarming such as the question we posed earlier concerning sensing of the environment by swarms.

**Figure 1.**
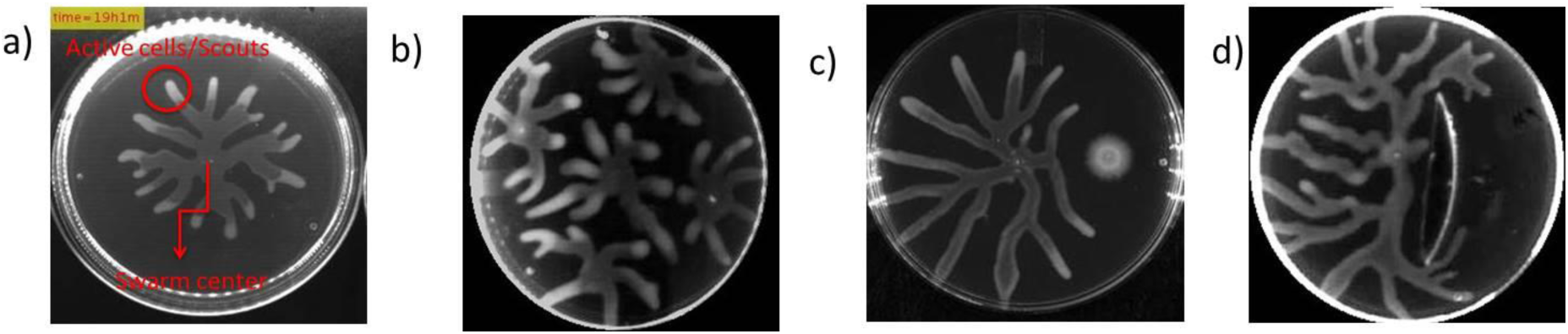
a) Representative labelled picture of a PA swarm, b) Avoidance behaviour among multiple colonies of wild type on the same petri-plate, c) Avoidance exhibited by the swarm colony of the wild type in the presence of flgM mutant, d) Avoidance exhibited by the swarm colony in the presence of a passive obstacle made of PDMS.

The tendrils produced by swarming PA are typically of the order of a centimetre with a width of 4-5 mm and their motion during swarming can be easily tracked with time-lapse imaging using regular digital cameras. This enables us to study the motion of the tendrils and their response to various environmental perturbations with relative ease compared to isotropically swarming bacteria (Kearns & Losick, 2003) and other bacterial species which form dense fractal-like patterns (Ben-Jacob, 1997). The motion of single bacterial cells on the agar surface can also be observed using GFP expressing bacteria leading to the generation of motility data spanning multiple spatial (microns to cm) and temporal (seconds to days) scales. Such multi-scale imaging data helps to link individual behaviour to the collective behaviour of the swarm. The observation and analysis of the response of the tendrils to perturbations introduced into the swarming medium (agar) reveals a remarkable ability of PA swarm to sense its spatial environment and respond to the presence of co-swarming sister tendrils (other tendrils from the same swarm initiating colony) [Figure 1 b)], approaching boundary of the petri-plate and even inert obstacles (Poly DiMethyl Siloxane (PDMS) and glass objects), several millimetres away along the swarming path. The ability to sense inert objects at millimetre scale distances is a particularly striking example of the spatial awareness of the PA swarm considering that the swarm is able to sense objects at distances three orders of magnitude (mm scale) further than body length of the individual members (micron scale). This fact is indicative of the collective advantage provided by the swarm as it is inconceivable that individual bacterium can sense inert objects at such long distances. We emphasize that the swarm is spatially aware of the object, i.e. the object does not actively secrete any molecules which are sensed by the bacterium.

The observation of spatial awareness by the swarm leads to the question of causative mechanisms. Using a continuous, fluid dynamic model involving secretion of a signalling molecule by the bacteria comprising the swarm, we show that a concentration gradient of the signalling molecule emerges within the swarm. This leads to two possible scenarios, namely, one in which the bacteria in different locations within the swarm behave differently leading to the collective response, or the other where the global gradient is implicitly computed by the swarm and to which it responds to. The latter scenario requires possibly complex information exchange within the swarm. To explore this further, we developed a multi-agent model, based on attractive-repulsive interaction arising purely from local information of the concentration of signalling molecules, which was able to replicate the spontaneous retraction (reversal of direction) of the swarm from an inert boundary. While this model is not yet a comprehensive representation of the swarming phenomenon, the ability to reproduce retraction suggests that the remarkable examples of spatial awareness seen in the PA swarm may not require active exchange of information between agents.

## Results

### General aspects of swarming pattern

The PA swarming system is interesting due to the highly specialized branching pattern [Figure 1 a)] and the remarkable spatial awareness and response of the swarm to its environment. A short survey of swarming patterns by *Paenibacillus dendritiformis, Paenibacillus vortex*, *Escherichia coli, Proteus mirabilis, Rhizobium etli, Serratia marcescens and Salmonella typhimurium* (Verstraeten et al., 2008)[SI Section 1 Figure S1] species of bacteria suggests that the PA swarming pattern occupies a unique parameter space relative to the other swarm patterns. Many bacterial species such as *E. coli* and *P. mirabilis* produce dense patterns which expand isotropically from the point of inoculation. *Bacillus subtilis* (Fujikawa and Matsushita 1989) *and Paenibacillus dendritiformis* under some conditions produce fractal like patterns which can be described by DLA models (Ben-Jacob, 1997). In contrast, PA swarms expand in straight segments (tendrils) from the inoculated region. The expansion of each tendril is highly anisotropic along a straight line with constant speed [SI Section 1 Figure S3]. The overall pattern is characterized by a robust statistical distribution of branch lengths and divergence angles [SI Section 1 Figure S2].

### Awareness of the Presence of Sister Tendrils

The swarming tendrils display self-avoidance (Caiazza et al. 2005)(Tremblay et al., 2007). This behaviour is also seen in growing *Bacillus subtilis* tendrils (James et al., 2009) and in P.. *dendritiformis* by Avraham Be’er et al (Be’er et al., 2009) who used the term sibling rivalry to describe this behaviour. In this case swarm fronts emerging from two locations in the same swarm plate lead to the formation of a zone of inhibition unpopulated by either of the advancing swarm fronts which can be viewed as a form of self-avoidance. However, there is a crucial difference between the self-avoidance seen in the PA system compared with the *Paenibacillus* system. As (Be’er et al., 2010) showed the zone of inhibition is due to mutually lethal secretions produced by the swarming *P. dendritiformis* resulting in the death of bacteria in the region between the advancing fronts producing the zone of inhibition. In the case of PA, the advancing sister tendrils sense each other and typically one of them retracts back (complete reversal) or changes its direction. [SI Section 2] This is a form of true self-avoidance and not a consequence of lethal secretions killing off the bacteria in the advancing front (James et al., 2009). In other words, the bacteria on one of the advancing tendrils sense the presence of another tendril advancing towards it and initiate changes in the motility profile which result in a change in the swarming direction. Thus, the self-avoidance phenomena in PA is significantly more complex and requires coordination far more in extent to that described previously in the case of *P. dendritiformis* and requires deeper study. We do not yet have a quantitatively accurate dynamical model of swarm direction reversal associated with the sensing of sister tendrils. However, the continuous model, based on sensing of the concentration gradient of a signalling molecule, presented later in the article is able to explain certain features such as the expected change in direction and why generally only one tendril retracts.

Another behaviour related to intra-species sensing is the avoidance of non-swarming mutant strains by advancing tendrils of swarming strains as shown in Figure 1 c) [SI Section 2 Figure S5 and video].The flgM mutant of PA is nonswarming due to the absence of flagella, yet it induces avoidance response in wild type PA. Thus this mutant likely indicates its presence, through secretions, sensed by the wildtype swarm which subsequently changes the direction of the advancing tendril.

### Awareness of the Presence of Petri-plate Boundary

As the tendrils approach the edge of the petri-plate, we often observe a complete reversal of swarming direction. Closer observation reveals that this effect happens most likely because one of the tendrils reaches the edge of the petri-plate and releases the swarming bacteria which move along the gap between the agar and the petri-plate wall. The signalling molecule(s) from these set of bacteria are sensed by the other tendrils which have not yet reached the edge and causes direction reversal in response to this sensory information. There is further support for this hypothesis as the advancing tendrils reverse their directions only after at least one of the tendrils has reached the edge of the petri-plate [SI Section 3 for video].

### Awareness of the Presence of Inert Objects

The examples of spatial awareness presented above are all related by the fact that the target for spatial awareness was of biological origin be it the tendrils of its own colony or that of a sister colony. The sensing mechanism in this case could be hypothesized to arise out of signalling molecules secreted by the targets. We investigated if the advancing tendrils could sense the presence of inert obstacles which would not secrete signalling molecules. Interestingly, we found strong evidence of the advancing tendril detecting the presence of the obstacle and changing its direction as it approached the obstacle. The bulk of our studies were conducted with the inert polymeric material PDMS. However, in order to check the material dependence of the obstacle, we also conducted more limited studies with obstacles made of glass [SI Section 4 Figure S6]. We did rigorous statistical analysis of experimental data to quantify the detection of inert obstacles by advancing swarm tendrils. We analysed around 120 experiments involving sensing of inert objects of various shapes with negative control (no obstacle) and positive control (flgM mutant) which causes the wild type strain to avoid it as described earlier. The positive control clearly shows that the swarming pattern is perturbed by the presence of the flagellar mutant. For the inert obstacle experiments we first evaluated the possibility that the perturbation seen in the swarming pattern indeed arises due to the presence of the obstacle. This was done by computationally scanning the obstacle shape across the image and finding the best fit position. A unique fit at the exact obstacle position or the best fit being around the obstacle position supports the hypothesis that the perturbation is due to the obstacle and not due to the natural stochasticity in the swarming pattern. This analysis revealed that about 80% of experiments, a significant majority, indicate successful detection of the inert obstacle by the swarm from as far as 5mm [see SI Section 4 for video]. The detection of obstacles was also observed with change in nutrient media (From PGM to M9) as with change of obstacle material. [See SI section 4 for videos]

In addition to this analysis, we quantified the asymmetry in the swarming pattern. This is motivated by the fact that the branch distribution in the case of a negative control can be assumed to be unbiased hence symmetric. However, the presence of an inert obstacle or the positive control creates asymmetry in the swarming pattern. To factor out the asymmetry due to the presence of the obstacles itself (as opposed to the perturbation due to the obstacle) we digitally replicated the image of the obstacle (inert or positive control) in all the four quadrants and subtract the obstacle region from the original image. The pattern obtained after subtracting the digitally created obstacles is used to calculate the variance of the swarm coverage in each quadrant. Both positive control and the experiments on inert obstacles showed significantly larger inter-quadrant variance indicating strong perturbation of the baseline swarming pattern. More details of the data analysis are provided in the SI text [Section 5]. The sensing of inert obstacles from a distance represents the strongest evidence of the advantages conferred by membership of the swarm as it is inconceivable that an individual bacterium can detect an inert object from such a long distance. In the subsequent section we propose a unified model to explain all these examples of spatial awareness in PA.

**Figure. 2.**
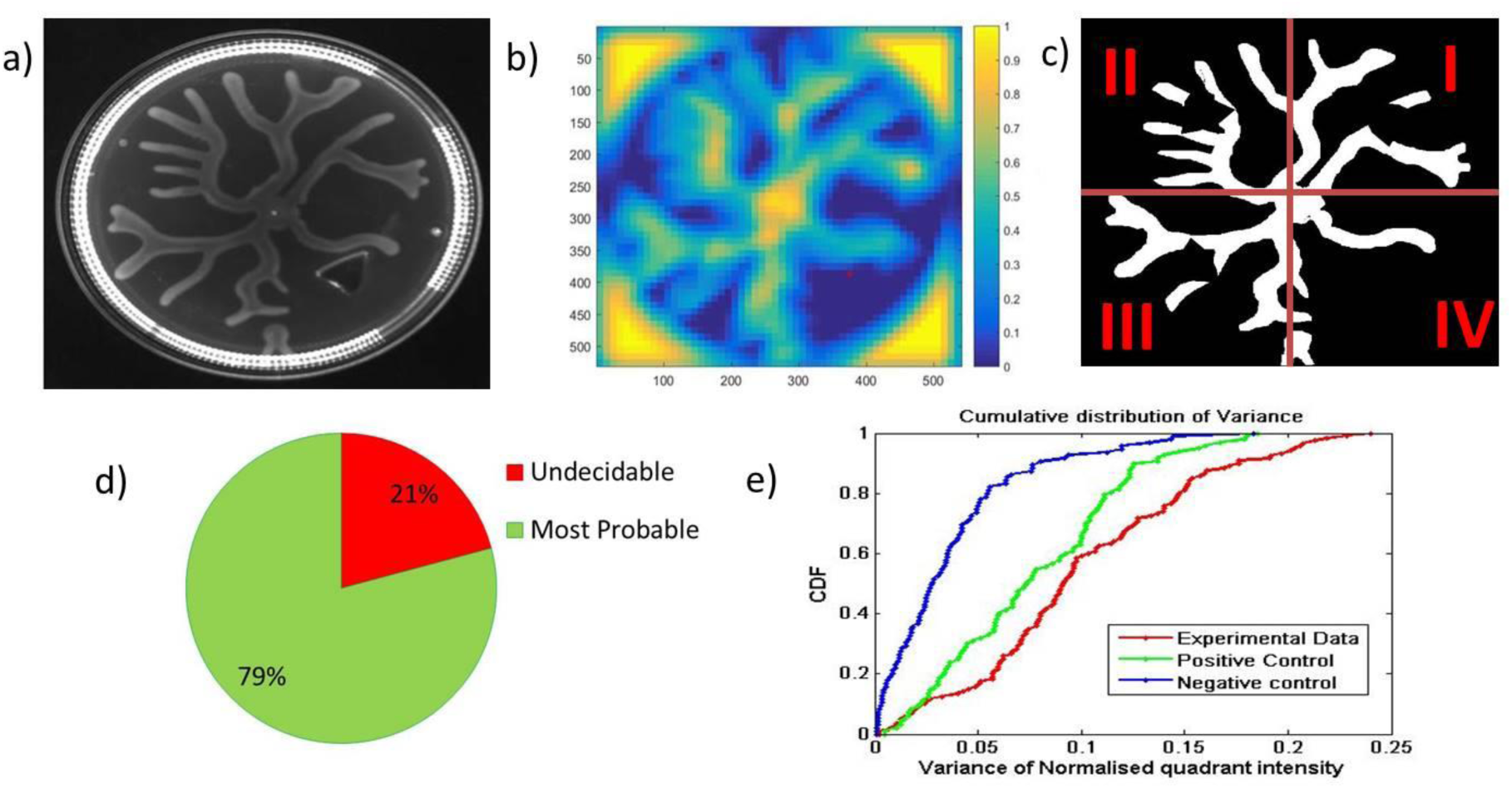
a) Swarm plate with PDMS obstacle, b) Heat map showing available positions of the given obstacle to occupy (blue: Obstacle will not overlap with the pattern, yellow: Obstacle overlaps with the pattern), c) Thresholded image pattern obtained by subtracting the obstacle for asymmetry analysis (See SI Section 5 for more details), d) Statistics of different types of perturbations to the pattern due to presence of the obstacle (Most probable: Patterns encircle the obstacle while avoiding it, Undecidable: Patterns have restricted growth and/or are unable to indicate the effect of the obstacle), e) Cumulative Distribution Function of inter-quadrant variance in the distribution of the area occupied by the swarming colony

### Towards a Mathematical Model of Spatial Awareness in the PA Swarm

The main idea of the mathematical model is that the individual bacterium and consequently the swarm can sense a signalling molecule (or a cocktail of molecules) to decide its direction. For the purpose of this model it is enough to consider a single molecule even though in practice several molecules may be involved. We perform quantitative modelling to understand a) coarse-grained dynamics of swarm direction changes induced by perturbations of the various kinds mentioned above and b) what specific advantages are available to the individual members of the swarm by virtue of membership in the swarm. The requirement of a signalling molecule leading to self-avoidance of sister tendrils as well as the other instances of spatial awareness mentioned in this article is well supported by previous reports describing the role of rhamnolipids (RL) in PA swarming [(Caiazza et al., 2005)(Tremblay et al., 2007)]. We suppose that the advancing swarm tip contains active (motile) bacteria which secrete signalling molecules at a specified rate f. The signalling molecules diffuse with diffusion constant D. The governing equations then become

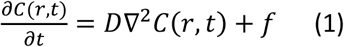

With f = 0 in the region outside the swarm tip. Inert obstacles or petri-plate edges are represented by reflecting (non-diffusive) boundaries. Parameters such as the production rate and diffusion constant of the signalling molecule, number of bacteria in a swarm are required to draw meaningful inferences from this model. Although these parameters have not been explicitly measured, we can estimate these from experiments if available, or by order of magnitude calculations [See SI Section 6 for more details]

Simulations proceed by initializing a circular disk of fixed radius representing the advancing swarm tip with bacteria secreting the signalling molecule. The concentration field of the signalling molecule is calculated using Eq. (1) above over the entire region and the “swarm tip” is advanced to a new position representing the motion of the swarming tendril. The concentration field is updated and the process continues. Firstly, we see that for the best estimates we have for the model parameters, a gradient in the order of µM/mm emerges within the swarm which is comparable to the gradients which bacterial species such as *E. coli* (Jeon et al., 2009)(Diao et al., 2006) have been reported to sense. We found that this gradient of the signalling molecule concentration accurately predicts the future direction of the swarm. Specifically, the swarm will move in the direction of the steepest negative gradient although inertia and stochastic effects would induce some deviations around this expected direction. These effects are not included in this model currently. However, this model serves to demonstrate that for reasonably realistic parameter values, measurable concentration differences of the signalling molecule emerge within the swarm which regulate the direction of the swarming tendril. In the case of sensing of sister tendrils, we see that µM/mm gradients appear typically only in the smaller swarm tip which then changes its direction of motion [Figure 3(a)(f)]. In the case of inert obstacles, we again see that the presence of a reflecting boundary results in measurable gradients which predict the future direction of the swarm [Figure 4]. For this case, we find the strongest argument for a bacterium’s requirement of membership in a swarm because measurable gradients from inert reflecting boundaries will only form if the source strength is large enough. The large source strength required is provided by the swarm whereas individual bacterium would never be able to produce it on its own.

**Figure. 3.**
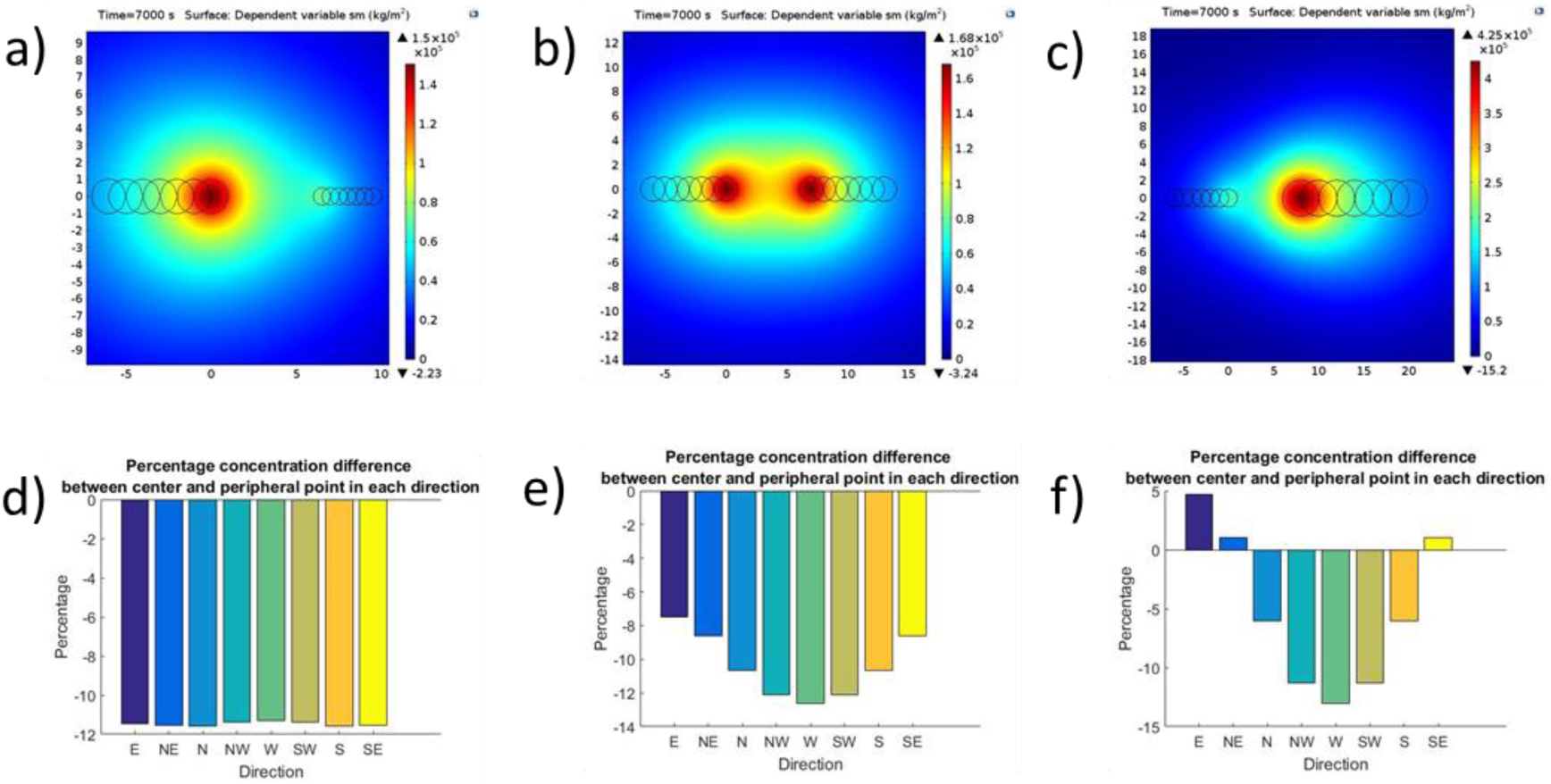
Signalling molecule gradient which emerges within the swarm tip when encountered by other approaching swarm tips of different sizes at a separation of 5mm.

**Figure 4.**
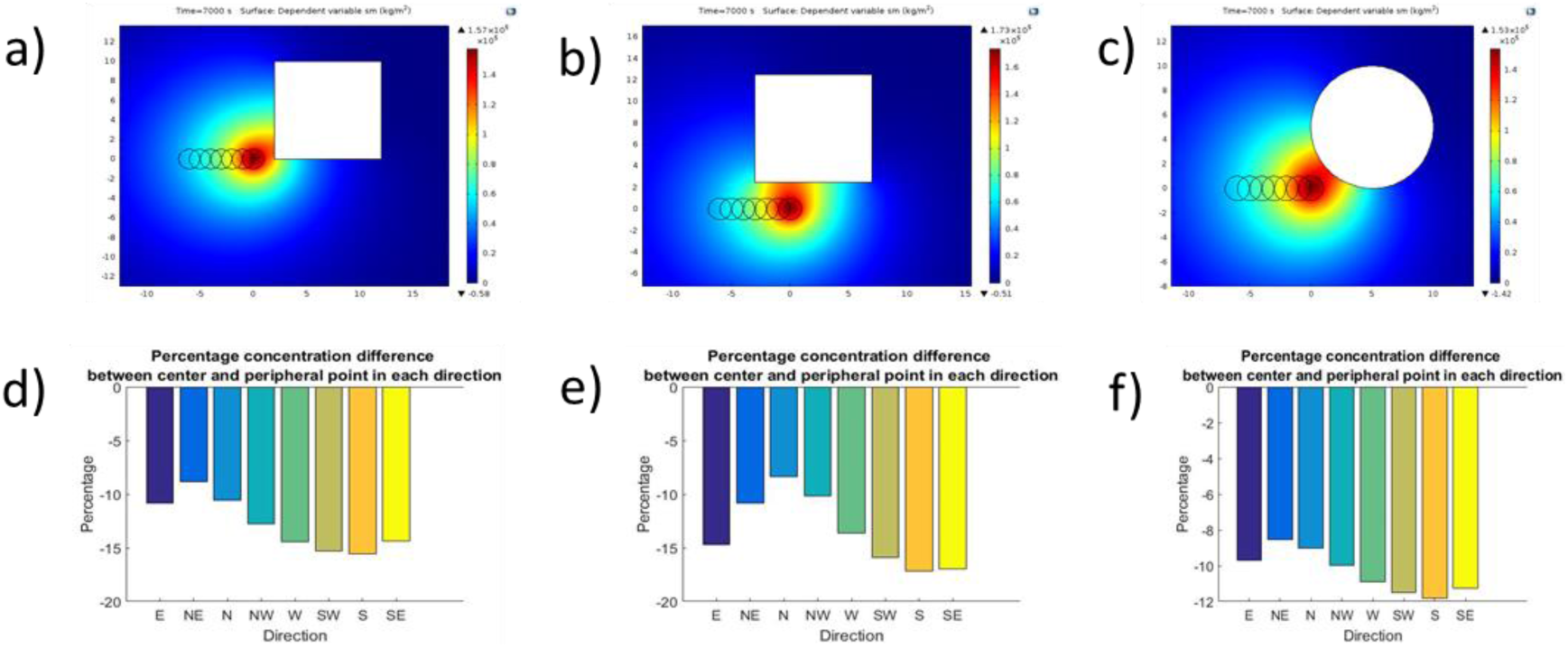
Signalling molecule gradient which emerges within the swarm tip due to an inert obstacle at a separation of around 1 mm.

The model discussed above is a coarse-grained model which assures us of the strong likelihood of measurable gradients of a signalling molecule forming within the swarm. Ultimately, we would like to understand collective spatial awareness of the swarm emerging from individual behaviour and cell-cell interactions including its inherent stochasticity. Insights derived from such studies can guide robust distributed control strategies for future robotic swarms (Rubenstein et al., 2012)] and other collective sensing phenomena. We constructed a multi-agent model [Wilensky, U. (1999). NetLogo] based on two-component signalling system. The two signalling molecules produced by the agents have different diffusion constants with each of them either invoking an attracting or repelling response among the agents. The spatial distribution of the signalling molecules govern the behaviour of the agent [See SI Section 8 for a detailed description of the multi-agent model], which spontaneously shows the emergence of branching as observed in experiments [Figure 5]. The agent based system also exhibited spontaneous emergence of spatial awareness similar to the biological system, most notably, the detection of boundaries (analogous to detection of inert obstacles presenting non-diffusive reflective boundaries) and consequent retraction of the advancing swarm tendril as observed in experiments. The spontaneous emergence of these features in the simulations suggest that the remarkable spatial awareness seen in the bacterial swarm may arise from simple attractive-repulsive interactions between bacteria which indirectly leads to the effective computation of the global gradient predicted by the fluid dynamic model.

**Figure. 5.**
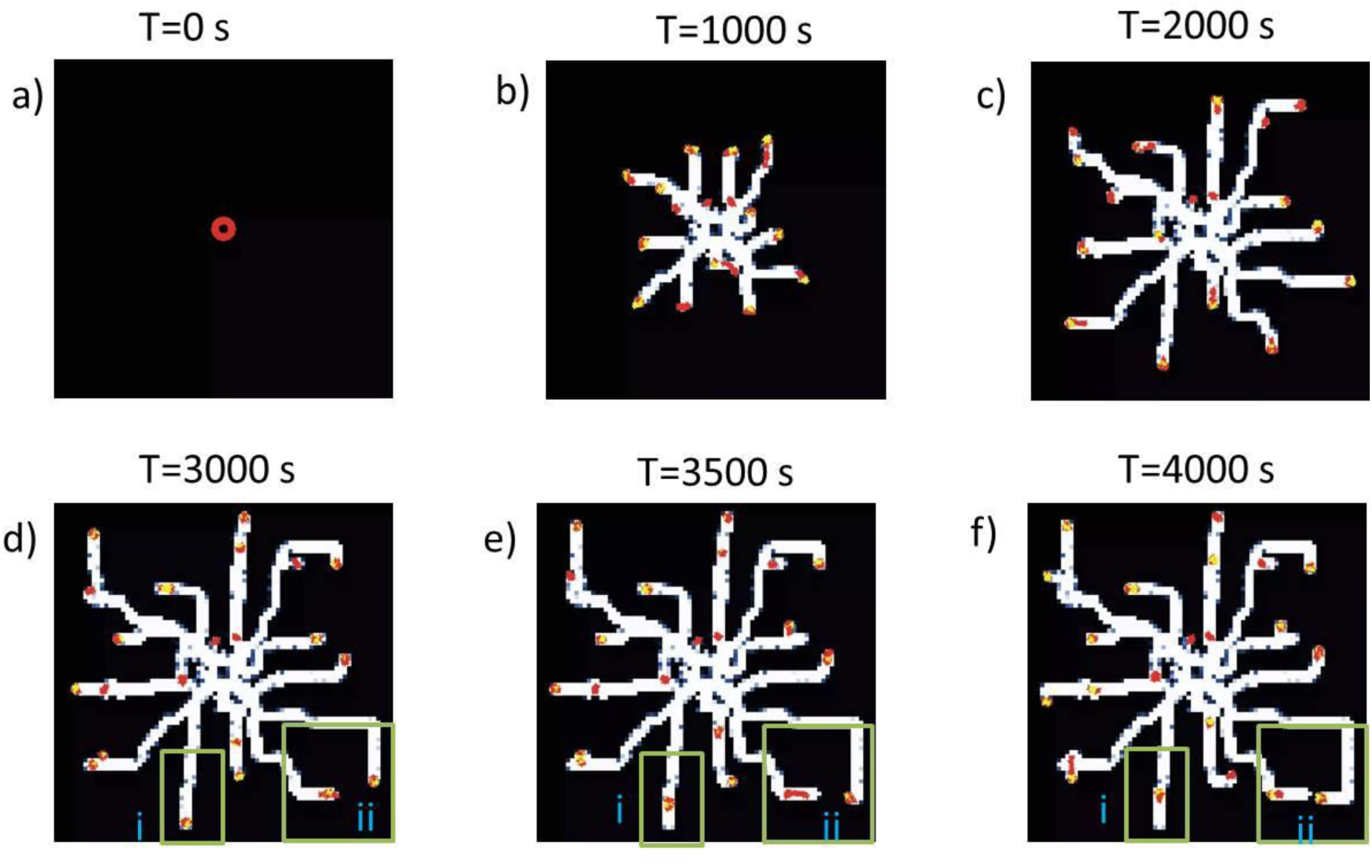
Illustration of agent based simulation showing branched pattern and retraction from boundary [Red pixels = Active swarmers in state 0, Yellow pixels = Active swarmers in state 1, White pixels = water film formed or colony spread (See SI section 7 for more details)]. Regions i and ii highlights the retraction phenomena from the boundary and other tendril respectively.

### Discussion and open questions

Detailed spatial awareness expectedly increases the fitness of an organism to survive a complex environment. In this article, we showed examples of a primitive organism displaying sophisticated spatial awareness such as long-range sensing of an inert obstacle. The response of the swarm to these perturbations is coherent and highly coordinated unlike previous descriptions of sensing neighbouring swarming colonies by lethal secretions. The role of a relatively fast diffusing signalling molecule is expected and strongly suggested by the mathematical models. In particular, the coarse-grain model provides strong evidence to the fact that measurable differences in the concentration of signalling molecules can arise from our expectations on the model parameters which will induce difference in behaviour in individual bacteria at the swarm boundaries. Indeed, a single cell resolved, multi-agent model based on signalling molecule concentration dependent attractive-repulsive potential exhibits spontaneous emergence of branching and some aspects of spatial awareness observed in the actual biological system. Presently the agreement between experiments and models is largely qualitative. The patterns observed in the multi-agent simulations do not resemble the patterns observed in the experiments indicating that the present model is not complete. Another open question is how the swarming tendril avoids self-inhibition from its own secretions while being able to inhibit the advancement of a neighbouring tendril at a much longer distance away. Although these questions are still open, this work presents some essential ingredients likely to lead to a quantitatively rigorous agent based model for PA swarming which can reproduce not only the pattern formation aspects but also its collective sensing abilities of the swarm. Such a model would also enable one to study robustness of the collective behaviour in the presence of defective individuals and other perturbations. Insights related to robust behaviour would be of exceptional value to the emerging field of swarm robotics from the perspective of robust decentralized control.

## Methods

### Swarming Motility Assay

For swarming assay, we used Peptone growth medium (PGM). Composition of PGM 0.6% agar plates are 6 grams of bacteriological agar (Bacto agar), 3.2 grams of peptone, and 3 grams of sodium chloride (NaCl) added in 1 litre of distilled water. The medium was autoclaved at 121°C for 30 minutes. After autoclaving the media, 1 mL of 1M CaCl2 (Calcium chloride), 1 mL of 1 M MgSO4 (Magnesium sulphate), 25 ml of 1M KPO4 and 1 mL of 5 mg/mL cholesterol were added into the medium and mixed properly. 25 mL PGM were poured in each 90mm Petri-plates and allowed them to solidify at room temperature (RT) for a half an hour under the laminar hood flow with the lid opened. And all the plates were kept at room temperature for 16-18 hours for further drying.

### Swarming

2ul of a planktonic culture of Wild type PA or flgM mutant with OD >2.8 is inoculated at the centre of 90 mm petri-plate containing PGM-0.6% agar. The wild type PA14 forms the pattern as shown in Figure 1 a) over a period of 24 hours in a 90 mm petri-plate. FlgM is a transposon insertion mutant in the flgM gene of PA14 and is part of P. aeruginosa transposon insertion library (Liberati et al., 2006).

### Preparation of PDMS (Poly DiMethyl Siloxane) obstacle

We have used Sylgard 184 from Dow Corning. It has two parts: an elastomer part and the curing agent. The two parts i.e. elastomer: curing agent is mixed in the ratio of 10:1. This mixture is stirred well. The air bubble cause due to stirring is removed by degassing the PDMS mixture in a desiccator connected to vacuum pump. An acrylic template is made to obtain different shapes of the obstacle. The air bubble free PDMS mixture is then poured into the template and cured for 12 hours. The cured PDMS solidifies and is removed from the acrylic template. These PDMS obstacles are then sterilised in autoclave.

The sterilised obstacle blocks are placed in an appropriate position in the petri-plate. The nutrient agar is then poured around the obstacle such that the obstacle is half immersed in the nutrient agar while held intact in its original position. The nutrient agar with the obstacle is allowed to dry under the laminar hood.

## Acknowledgements

We gratefully acknowledge Robert Bosch Centre for Cyber Physical Systems at Indian institute of Science, Bangalore, India for funding this research. We also acknowledge the use of facilities at Centre for Nano Science and Engineering, Indian Institute of Science, Bangalore, India.

## Reference

Be’er, A., Ariel, G., Kalisman, O., Helman, Y., Sirota-Madi, A., Zhang, H. P., … Swinney, H. L. (2010). Lethal protein produced in response to competition between sibling bacterial colonies. Proceedings of the National Academy of Sciences of the United States of America, 107(14), 6258–6263. http://doi.org/10.1073/pnas.1001062107

Be’er, A., Zhang, H. P., Florin, E.-L., Payne, S. M., Ben-Jacob, E., & Swinney, H. L. (2009). Deadly competition between sibling bacterial colonies. Proceedings of the National Academy of Sciences of the United States of America, 106(2), 428–433. http://doi.org/10.1073/pnas.0811816106

Ben-Jacob, E. (1997). From snowflake formation to growth of bacterial colonies II: Cooperative formation of complex colonial patterns. Contemporary Physics, 38(3), 205–241. http://doi.org/10.1080/001075197182405

Caiazza, N. C., Shanks, R. M. Q., & Toole, G. A. O. (2005). Rhamnolipids Modulate Swarming Motility Patterns of Pseudomonas aeruginosa. Journal of Bacteriology, 187(21), 7351–7361. http://doi.org/10.1128/JB.187.21.7351–7361.2005

Daniels, R., Vanderleyden, J., & Michiels, J. (2004). Quorum sensing and swarming migration in bacteria. FEMS Microbiology Reviews, 28, 261–289. http://doi.org/10.1016/j.femsre.2003.09.004

Deng, P., Roditi, L. D. V., Ditmarsch, D. Van, & Xavier, J. B. (2014). The ecological basis of morphogenesis: branching patterns in swarming colonies of bacteria. New Journal of Physics. http://doi.org/10.1088/1367-2630/16/1/015006

Diao, J., Young, L., Kim, S., Fogarty, E. a, Heilman, S. M., Zhou, P., … DeLisa, M. P. (2006). A three-channel microfluidic device for generating static linear gradients and its application to the quantitative analysis of bacterial chemotaxis. Lab on a Chip, 6, 381–388. http://doi.org/10.1039/b511958h

Du, H., Xu, Z., Anyan, M., Kim, O., Leevy, W. M., Shrout, J. D., & Alber, M. (2012). High density waves of the bacterium pseudomonas aeruginosa in propagating swarms result in efficient colonization of surfaces. Biophysical Journal, 103, 601–609. http://doi.org/10.1016/j.bpj.2012.06.035

Du, H., Xu, Z., Shrout, J. D., & Alber, M. (2011). MULTISCALE MODELING OF PSEUDOMONAS AERUGINOSA SWARMING. Mathematical Models and Methods in Applied Sciences, M3AS(21 Suppl 1), 939–954. http://doi.org/10.1142/S0218202511005428

James, B. L., Kret, J., Patrick, J. E., Kearns, D. B., & Fall, R. (2009). Growing Bacillus subtilis tendrils sense and avoid each other: Research letter. FEMS Microbiology Letters, 298, 12–19. http://doi.org/10.1111/j.1574-6968.2009.01665.x

Jeon, H., Lee, Y., Jin, S., Koo, S., Lee, C. S., & Yoo, J. Y. (2009). Quantitative analysis of single bacterial chemotaxis using a linear concentration gradient microchannel. Biomedical Microdevices, 11, 1135–1143. http://doi.org/10.1007/s10544-009-9330-8

Kearns, D. B. (2010). A field guide to bacterial swarming motility. Nature Reviews Microbiology, 8, 634–644. http://doi.org/10.1038/nrmicro2405

Kearns, D. B., & Losick, R. (2003). Swarming motility in undomesticated Bacillus subtilis. Molecular Microbiology, 49(3), 581–590. http://doi.org/10.1046/j.1365-2958.2003.03584.x

Liberati, N. T., Urbach, J. M., Miyata, S., Lee, D. G., Drenkard, E., Wu, G., … Ausubel, F. M. (2006). An ordered, nonredundant library of Pseudomonas aeruginosa strain PA14 transposon insertion mutants. Proceedings of the National Academy of Sciences of the United States of America, 103, 2833–2838. http://doi.org/10.1073/pnas.0511100103

Rubenstein, M., Ahler, C., & Nagpal, R. (2012). Kilobot: A low cost scalable robot system for collective behaviors. Proceedings - IEEE International Conference on Robotics and Automation, 3293–3298. http://doi.org/10.1109/ICRA.2012.6224638

Tremblay, J., Richardson, A., Lépine, F., & Déziel, E. (2007). Self-produced extracellular stimuli modulate the Pseudomonas aeruginosa swarming motility behaviour. Environmental Microbiology, 9(10), 2622–2630. http://doi.org/10.1111/j.1462-2920.2007.01396.x

Verstraeten, N., Braeken, K., Debkumari, B., Fauvart, M., Fransaer, J., Vermant, J., & Michiels, J. (2008). Living on a surface: swarming and biofilm formation. Trends in Microbiology, 16(10), 496–506. http://doi.org/10.1016/j.tim.2008.07.004

Vicsek, T., Czirok, A., Ben-Jacob, E., Cohen, I., & Shochet, O. (1995). Novel Type of Phase Transition in a System of Self-Driven Particles. Physical Review Letters, 75(6), 1226–1229.

